# Precursor abundance influences divergent antigen specific CD8^+^ T cell responses after *Yersinia pseudotuberculosis* foodborne infection

**DOI:** 10.1101/2021.05.10.443536

**Authors:** Yue Zhang, Zhijuan Qiu, Brian S. Sheridan, James B. Bliska

## Abstract

Primary infection of C57BL/6 mice with the bacterial pathogen *Yersinia pseudotuberculosis* elicits an unusually large H-2K^b^-restricted CD8^+^ T cell response to the endogenous and protective bacterial epitope YopE_69-77_. To better understand the basis for this large response, the model OVA_257-264_ epitope was inserted into YopE in *Y. pseudotuberculosis* and antigen specific CD8^+^ T cells in mice were characterized after foodborne infection with the resulting strain. The epitope YopE_69-77_ elicited significantly larger CD8^+^ T cell populations in the small intestine, mesenteric lymph nodes (MLNs), spleen and liver between 7- and 30-days post infection, despite residing in the same protein and having a similar affinity for H-2K^b^ as OVA_257-264_. YopE-specific CD8^+^ T cell precursors were ~4.6 times as abundant as OVA-specific precursors in the MLNs, spleen and other lymph nodes of naïve mice, explaining the dominance of YopE_69-77_ over OVA_257-264_ at early infection times. However, other factors contributed to this dominance as the ratio of YopE-specific to OVA-specific CD8^+^ T cells increased between 7- and 30-days post infection. We also compared the YopE-specific and OVA-specific CD8^+^ T cells generated during infection for effector and memory phenotypes. Significantly higher percentages of YopE-specific cells were characterized as short short-lived effectors while higher percentages of OVA-specific cells were memory precursor effectors at day 30 post infection in spleen and liver. Our results suggest that a large precursor number contributes to the dominance and effector and memory functions of CD8^+^ T cells generated to the protective YopE_69-77_ epitope during *Y. pseudotuberculosis* infection of C57BL/6 mice.

## Introduction

CD8^+^ T cells are vital for defense against intracellular pathogens and malignant diseases. They are responsible for eliminating altered cells--those infected with intracellular microbes or tumor cells. Antigen-specific CD8^+^ T cells recognize peptide antigens presented on the Major Histocompatibility Class I (MHC I) molecules on the surface of these altered cells. Activated CD8^+^ cells then kill target cells through secretion of cytokines or expression of effector molecules (1). Identification of factors regulating the antigen specific CD8^+^ T cell response is critical for vaccine design to combat infectious diseases. In addition, bacteria have recently been recognized as a potential vaccine vector to stimulate CD8^+^ T cell response to treat cancer (2). CD8^+^ T cell response to bacteria is most thoroughly studied using Gram-positive pathogen *Listeria monocytogenes* (Lm) (3). Primary CD8^+^ T cell response to Lm can be divided into four phases--activation, expansion, contraction, and memory (3, 4). Following Lm infection, antigen-presenting cells (APCs), most notably Batf3-dependent CD8α^+^ dendritic cells (DCs) are required to acquire and present antigens to specific CD8^+^ T cell precursors in the T cell zone in the secondary lymphoid organs (5, 6). Antigen specific precursors are very rare, yet the process of antigen presentation to the precursors is surprisingly fast and efficient (7). Activated antigen-specific CD8^+^ T cells then undergo rapid expansion, a process requiring co-stimulatory factors and inflammatory cytokines including interleukin (IL)-2 and IL-12 (8, 9). Depending on the nature of the inflammatory environment, one activated precursor CD8^+^ T cell can give rise to more than 10,000 daughter cells over the next 5-8 days (10–14). Such rapid proliferation was estimated to be near the maximal possible division speed for mammalian cells. This expansion phase ensures the fastest accumulation of CD8^+^ T cells to accomplish their tasks (1). Coupled to this rapid expansion phase, the daughter cells differentiate into either short-lived effector cells or memory precursor effector cells. Local environments including inflammatory and antigenic stimuli play an important role in deciding the differentiation path of the daughter cells (14–16). For acute infection, after the rapid proliferation stage and regardless of whether the T cell response is successful in eliminating the pathogen or not, more than 90-95% of the expanded cells undergo apoptosis to result in contraction of the population (10, 17). Inflammation and especially the inflammatory cytokines interferon γ (IFNγ) and IL-12 influence the contraction phase (4). In addition, IL-7, IL-15 and transforming growth factor β (TGFβ) have all been implicated in the process (3). In the end, the remaining long-lived memory population provides faster and enhanced protection for secondary antigenic challenge (3, 4). Among the memory T cells generated, the tissue-resident memory (T_RM_) cells provide immediate local protection against subsequent infections (15).

The gram-negative bacterial pathogen *Yersinia pseudotuberculosis* has become a new model system to study CD8^+^ T cell response to bacteria. As an enteric zoonotic pathogen, *Y. pseudotuberculosis* is most commonly associated with mesenteric lymphadenitis in humans (18). Experimental infection of mice through the oral route with *Y. pseudotuberculosis* results in bacterial colonization in Peyer’s patches, the lamina propria, and mesenteric lymph nodes (MLNs). Bacteria also disseminate to the spleen and liver (19, 20). Previously, we showed that sub-lethal infection of C57BL/6 mice with *Y. pseudotuberculosis* resulted in a large H-2K^b^-restricted CD8^+^ T cell response to YopE_69-77_, which is a protective epitope found in one of the major virulence factors of *Yersinia* (21–26). *Y. pseudotuberculosis* is an extracellular pathogen and most of the bacteria have been found extracellularly *in vivo* during infection, though a small proportion of intracellular bacteria do survive at least temporarily (27–29). This extracellular localization of the bacteria is due to the effects of several anti-phagocytic proteins of *Yersinia*, including YopE. These proteins are translocated into the cytosol of host cells through the bacterial type III secretion system (T3SS) (30). T3SS is utilized by several Gram-negative pathogens to promote virulence (31). In *Yersinia*, activation of the T3SS requires close contact between a bacterium and the host cell (32). Evidence suggests that YopE-specific CD8^+^ T cells participate in host protection against *Yersinia* by the production of TNFα and IFNγ (22). Alternatively, it was suggested that CD8^+^ T cells use CTL activity to kill host cells attached to *Yersinia*, followed by engulfment and destruction of both the host cell and the attached bacteria by neighboring phagocytes (33). The dominant YopE-specific CD8^+^ T cell response requires CCR2^+^ inflammatory monocytes, which are recruited to foci of *Y. pseudotuberculosis* infection in lymphoid tissues (25, 26). These monocytes, and presumably their differentiated products such as inflammatory dendritic cells (DCs), are injected with YopE and are required to function as APCs to directly prime YopE-specific CD8^+^ T cells (25, 26).

To further characterize the CD8^+^ T cell response to *Y. pseudotuberculosis*, the model CD8^+^ T cell epitope in chicken ovalbumin (OVA_257-264_) was chosen to take advantage of existing tools, and because the OVA antigen has a similar predicted affinity for H-2K^b^ as YopE_69-77_ (24). We generated a *Y. pseudotuberculosis* strain in which the OVA_257-264_ sequence was inserted within a linker region of the native YopE protein, ensuring that both epitopes are equally produced during infection. Following foodborne infection of C57BL/6 mice with this strain, we unexpectedly found that the YopE-specific CD8^+^ T cell response was several times higher than that of the OVA-specific response in all tissues analyzed and at different days post infection. Furthermore, in naïve mice, YopE-specific CD8^+^ T cell precursors were ~4.6 times as abundant as the OVA-specific cells. In addition, YopE-specific and OVA-specific CD8^+^ T cells displayed differences in effector and memory formation. Higher percentages of YopE-specific cells were characterized as short-lived effector cells while higher percentages of OVA-specific cells were memory precursor effector cells. Together, these results support the proposition that the precursor frequency of antigen specific CD8^+^ T cells contribute to dominance during the primary immune response and may impact effector and memory functions in foodborne *Y. pseudotuberculosis* infection.

## Results

### Characterization of 32777-OVA for tissue colonization after foodborne infection

Previously Bergsbaken et al. measured CD8^+^ T cell responses to OVA_257-264_ and YopE_69-77_ in mice that were infected with a *Y. pseudotuberculosis* strain that expressed both native YopE and a YopE-ovalbumin fusion protein under a heterologous promoter (37). In that case, 4-5 times more YopE-specific than OVA-specific CD8^+^ T cells were produced in various tissues at the peak (day 9) and memory (day 45) phases of the response (37). Bergsbaken et al. reasoned that the higher antigen load of YopE_69-77_ as compared to OVA_257-264_ resulted in this difference in antigen-specific CD8^+^ T cell response, since ovalbumin was produced at lower levels relative to that of YopE (37). Here, to ensure equal antigen load, a *Y. pseudotuberculosis* strain 32777-OVA was generated by inserting the SIINFEKL (OVA_257-264_) epitope into a linker region of the YopE protein between the chaperone-binding domain and the GAP domain (Fig. 1A). This placed OVA_257-264_ adjacent to YopE_69-77_, which was predicted to possess a comparable affinity for MHC-I H-2K^b^ (IC50 of 17 nM and 20 nM, respectively) (24). C57BL/6 mice were infected by the foodborne route with a sublethal dose of 32777-OVA to characterize tissue colonization. From 4-14 days post infection (dpi), mesenteric lymph nodes (MLNs), livers and spleens were found to be colonized at variable levels. In general, the mean colonization levels decreased gradually between 7 and 14 dpi and were cleared at 30 dpi (Fig. 1 BCD). Similar to what we observed before for the parental wild-type strain 32777 (38), colonization by 32777-OVA ranged greatly from individual to individual, regardless of the tissues analyzed. For example, in spleens at 9 dpi, even from the same experiment, colonization ranged from below detection to a log of 8 (Fig. 1D). At 14 dpi, when the majority of the animals cleared bacteria from this site, two mice harbored moderate CFU levels in spleens (Fig. 1D). It is possible that insertion of the OVA_257-264_ sequence in YopE resulted in attenuation to some degree; however, we reasoned that tissue colonization by 32777-OVA was sufficient to elicit OVA-specific and YopE-specific CD8^+^ T cells after foodborne infection (see below).

**Fig. 1.**
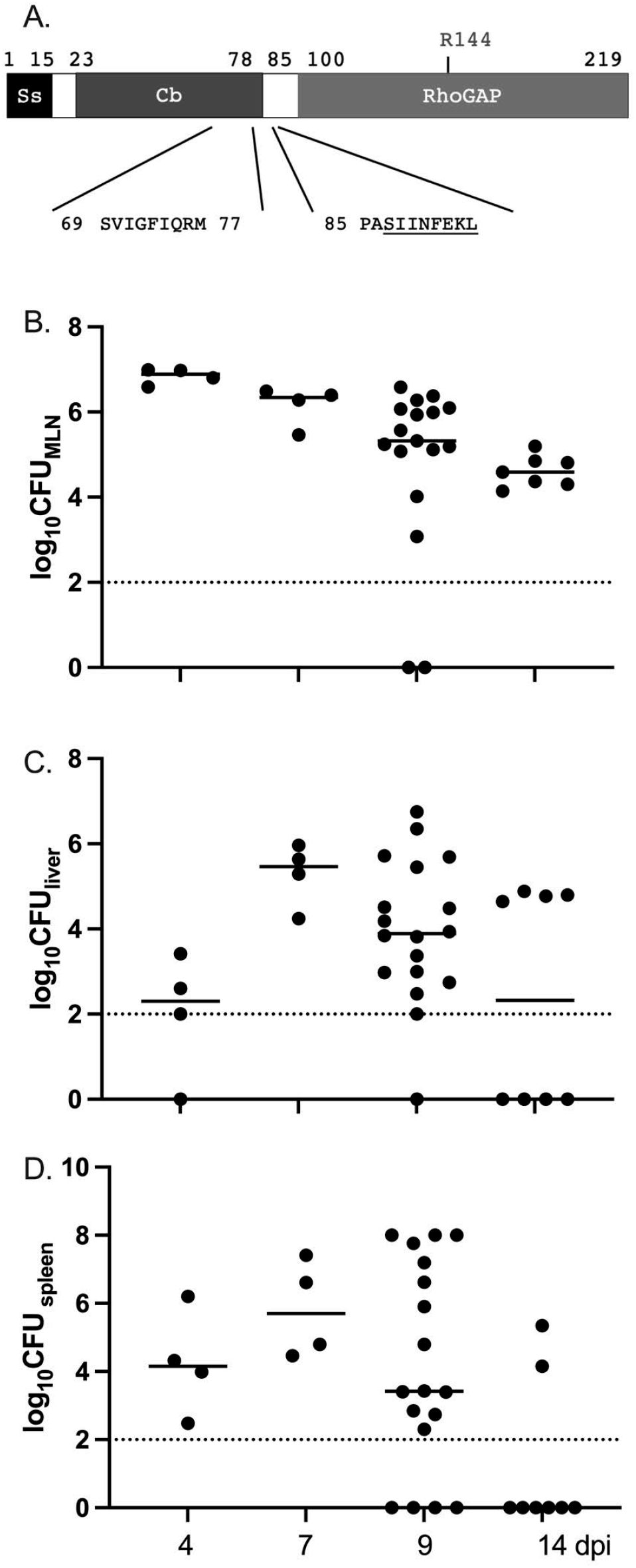
Structure of YopE protein in 32777-OVA and the tissue colonization levels of 32777-OVA following foodborne infection. (A), The domain structure of YopE protein, with signal sequence (Ss) for its secretion through the type 3 secretion system, the chaperone-binding domain (Cb) and the RhoGAP domain indicated. The arginine residue critical for the RhoGAP function is also indicated. The sequence and position of YopE_69-77_ and the OVA_257-264_ epitope inserted after residue 85 are indicated below. (B-D), C57BL/6 mice were infected with 5X10^7^ CFU of 32777-OVA orally and the colonization levels were determined by CFU assay at the indicated days post infection (dpi) from MLN (B), liver (C) and spleen (D). Each point represents the value obtained from one mouse and the results shown are pooled from 2-6 independent experiments, with *n* equals 4 for 4 dpi and 7 dpi, 18 for 9 dpi, 7 for MLN at 14 dpi, and the rest 8 for 14 dpi and 30 dpi. Dotted line indicates limit of detection. Means are indicated with a bar.

### Higher levels of YopE-specific compared to OVA-specific CD8^+^ T cells after 32777-OVA infection

With 32777-OVA foodborne infection, YopE-specific CD8^+^ T cells could be detected above background levels using tetramers and flow cytometry starting at 6 dpi (data not shown) and by 7 dpi a sizable population was evident in all the tissues analyzed—MLN, liver, and spleen (Fig. 2A). Throughout the time course of the analysis from 7-30 dpi, because bacterial colonization impacts inflammatory cytokines and correlates with antigen load, and there was a wide range in bacterial colonization levels from mouse to mouse, the levels of YopE-tetramer positive cells ranged widely among the individual animals analyzed (Fig. 2B-E). In general, on average the percentages of YopE-specific CD8^+^ T cells in each tissue increased between 7 and 9 dpi and then remained high over the remaining time course. In different tissues, the number of YopE-specific CD8^+^ T cells peaked at different days post infection, with spleens harbored the largest numbers of these cells while livers contained the highest percentage (Fig. 2C-E). Unexpectedly, in all tissues analyzed at all the time points, a smaller percentage of OVA-specific CD8^+^ T cells were identified (Fig. 2A-E). Furthermore, the percentage of these cells increased at a smaller magnitude from 7 to 9 dpi. As a result, the ratios of the percentages of YopE-specific/OVA-specific CD8^+^ T cells in MLNs, livers and spleens increased gradually from 7 to 30 dpi, and in all these tissues, the differences between 7 dpi and 30 dpi were significant (Fig. 2 F-H). In conclusion, in all tissues analyzed at all times, there were several times as many YopE-specific CD8^+^ T cells as OVA-specific cells; and the difference between the two populations increased from 7 to 30 dpi.

**Fig. 2.**
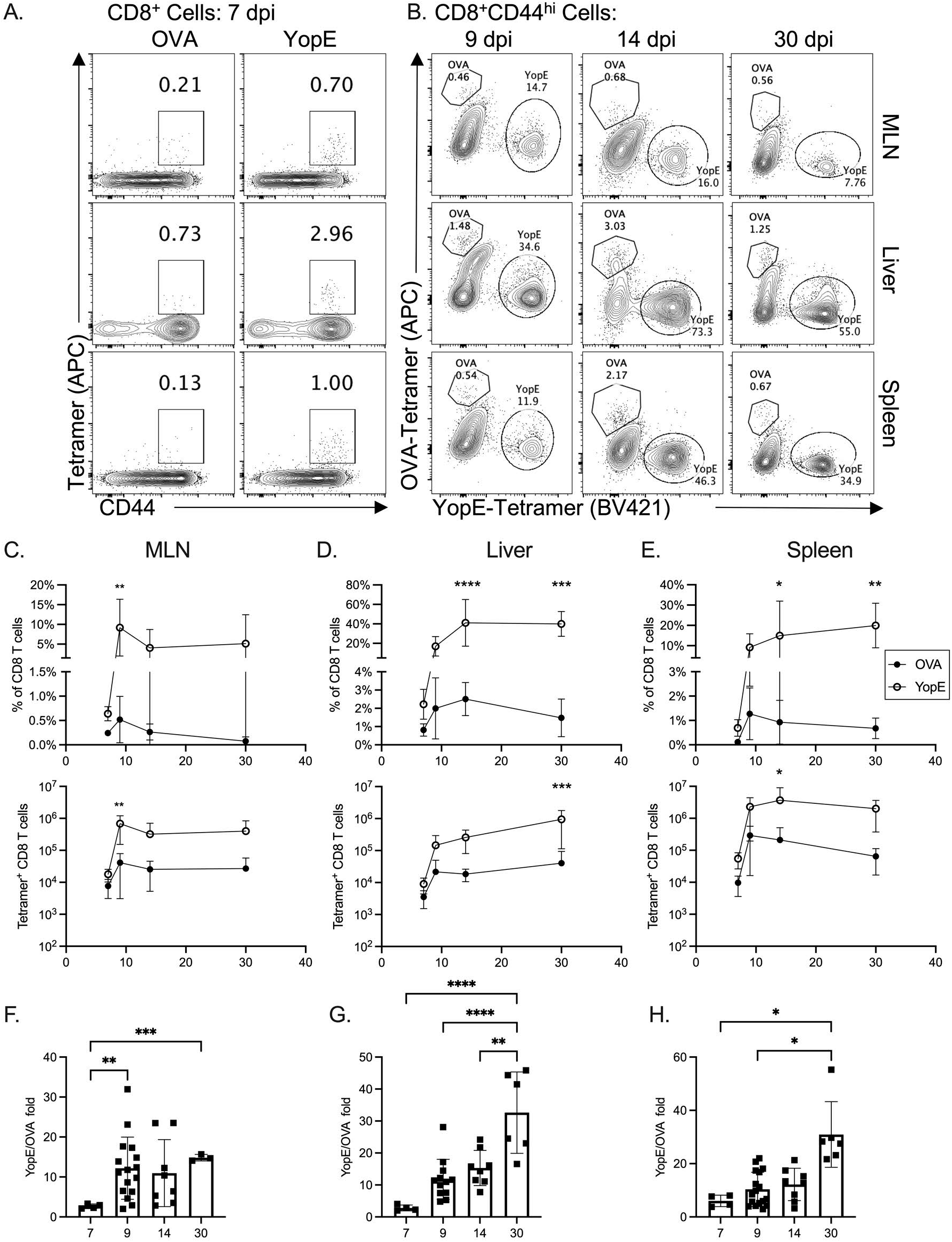
Comparison of the levels of OVA-specific and YopE-specific CD8^+^ T cell over time after infection with 32777-OVA. C57BL/6 mice were orally infected as in Fig. 1 with 32777-OVA. At 7, 9, 14 or 30 dpi, single cell suspensions from indicated tissues were prepared and analyzed with flow cytometry after staining with tetramers and antibodies. A., Representative contour plots of gated alive CD45.2^+^CD3^+^CD8^+^ T cells at 7 dpi from MLN (top), liver (middle) and spleen (bottom) further analyzed for CD44 and OVA-specific- (left) or YopE-specific- (right) signals. B. Representative contour plots of alive CD45^+^TCRβ^+^CD8α^+^CD44^hi^ cells from the indicated tissues at 9, 14 or 30 dpi analyzed for OVA- and YopE-tetramer expression. The percentages (top) and number (bottom) of alive tetramer-positive CD8^+^ T cells from MLNs (C), livers (D), and spleens (E) of mice infected for the indicated days. Filled circles indicate that of OVA-specific cells and open circle YopE-specific ones. Each point represents the value obtained from one mouse and the results shown are pooled from 2 independent experiments. For 7 dpi, *n* equals 4; 9 dpi, 7; 14 dpi, 8 and 30 dpi, 6. Means and standard deviations are shown. P values were determined with Two-way ANOVA (Mixed-effects model) followed by Sidak’s multiple comparisons test and values smaller than 0.05 are indicated. (F-H), the ratio of the percentages of YopE-specific CD8^+^ cells over that of OVA-specific ones at indicated dpi. The samples that no OVA-specific cells were detected or the percentage of OVA-specific ones were less than 0.01% were removed from analysis. F., the ratio of YopE/OVA from MLNs, with *n* equals 4, 16, 8 and 3, respectively, for 7, 9, 14 and 30 dpi. G., ratio of YopE/OVA from livers, with *n* equals 4, 12, 8 and 6 respectively for 7, 9, 14 and 30 dpi. H., ratio of YopE/OVA from spleens, with *n* equals 4, 17, 8, and 6 respectively for 7, 9, 14 and 30 dpi. Results shown are summary from 2-5 experiments. P values were determined with Brown-Forsythe ANOVA test followed with Dunnett’s T3 multiple comparisons test and values smaller than 0.05 are indicated.

### Higher levels of YopE-specific compared to OVA-specific CD8^+^ T cell precursors before 32777-OVA infection

The number of endogenous CD8^+^ T cell precursors of distinct specificities is known to be different, and the precursor frequency has been indicated to influence the size and progress of both the initial and the memory immune response (39). We consistently detected higher numbers of YopE-specific as compared to OVA-specific CD8^+^ T cells at the earliest infection times, raising the possibility that there is a difference in precursor numbers. Therefore, the numbers of YopE- and OVA-specific CD8^+^ T cell precursors in C57BL/6 mice was determined (Fig. 3ABC). Cells from the spleen, MLN and other macroscopically identifiable lymph nodes including inguinal, axillary, brachial, submandibular, cervical and para-aortic nodes were pooled from individual naïve mice and precursors were enumerated using the established protocol of peptide-MHC-I (pMHC) tetramer staining in conjunction with magnetic-bead separation (39). The average number of OVA-specific CD8^+^ T cells per mouse was determined to be 302 (Fig. 3AC), which is higher than what has been measured initially as ~130 (39), but is in range with other measurements reported later on (40). In contrast, the average number of YopE-specific CD8^+^ T cells was determined to be 1387 per mouse, or 4.6 times the number measured for OVA-specific cells (Fig. 3BC). Thus, there are significantly more YopE-specific CD8^+^ T precursors than OVA-specific cells in C57BL/6 mice, which likely contributes to the dominance of the former.

**Fig. 3.**
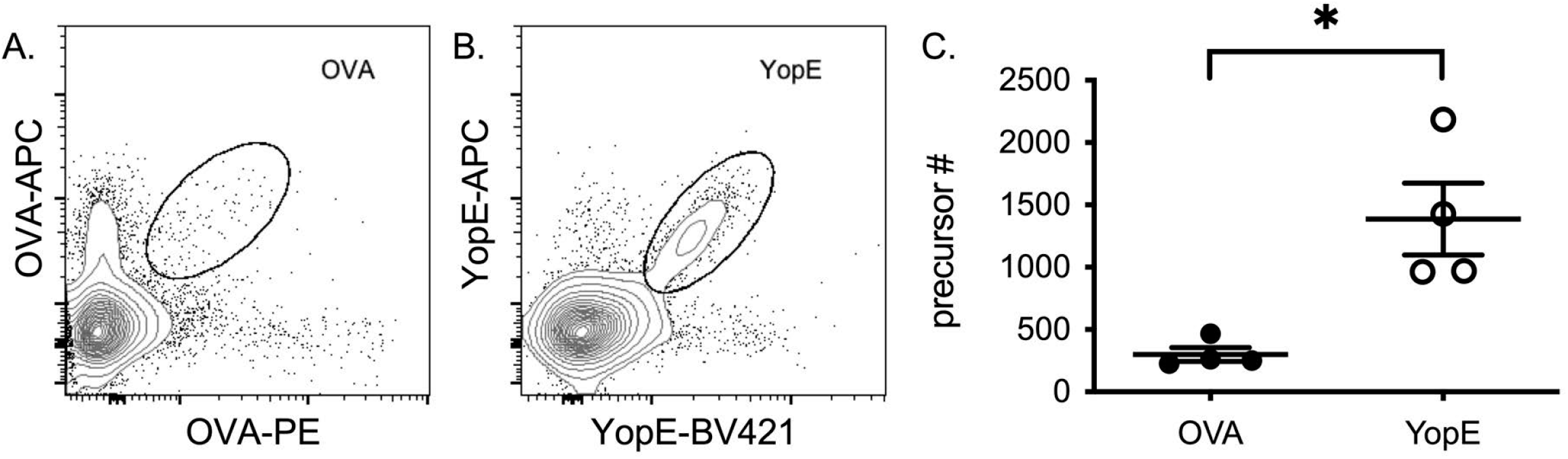
Precursor determination of OVA- and YopE-specific CD8^+^ T cells in naïve C57BL/6 mice. Representative dot plots of the enriched OVA (A) or YopE (B) tetramer-positive cells as indicated. (C), The absolute numbers of the naïve antigen specific CD8^+^ T cells from total spleen and lymph nodes of individual naïve mice. Each symbol represents the value obtained from an individual mouse <u>where *n* equals 4</u> and mean values and standard deviations are indicated. P value was determined by Mann-Whitney test.

### Characterization of YopE-specific and OVA-specific CD8^+^ T cell effector phenotypes generated during 32777-OVA infection

We next determined if the dominance of YopE-specific over OVA-specific CD8^+^ T cells impacted their effector functions. <u>Initially</u>, the effector phenotype of YopE- and OVA-specific CD8^+^ T cells were evaluated through accessing the expression levels of the terminal effector marker KLRG1 at 9 dpi. As expected, the levels of KLRG1^high^ cells varied among the tissues analyzed, and in general, blood contained the highest levels of KLRG1^high^ antigen specific CD8^+^ cells and the percentages of KLRG1^high^ antigen-specific cells were lower in MLN, liver and spleen (Fig. 4). Curiously, the YopE-specific CD8^+^ T cells contained lower levels of KLRG^high^ cells than the OVA-specific ones. In blood, an average of 83% of OVA-specific CD8^+^ T cells were KLRG^high^, while in contrast, only an average of 59% of YopE-specific ones were KLRG^high^ (Fig. 4A, top panels and B, first two columns). Significantly lower levels of KLRG^high^ YopE-specific CD8^+^ T cells were also observed in the spleen and liver (Fig. 4AB). The levels of KLRG^high^ YopE-specific CD8^+^ T cells also trended lower in MLN, but the difference was not significant (P=0.06, Fig. 4B). Overall, the difference in the levels of KLRG1^high^ cells suggests a potential difference in the differentiation of YopE-specific and OVA-specific CD8^+^ T cells at 9dpi. Next, these antigen specific CD8^+^ T cells at 9dpi were assessed through measuring the percentages that produced IFNγ or TNFα in response to peptide stimulation. Aliquots of cells from MLN or spleen of mice infected with 32777-OVA 9 days previously were stimulated *ex vivo* for 4.5 h with either YopE or OVA peptides or left non-stimulated (NS) in the presence of BFA, followed by intracellular cytokine staining and flow cytometry analysis (Fig. 5A). As expected from the dominant presence of YopE-specific CD8^+^ T cells, in comparison to cells treated with OVA peptide, or NS, a higher percentage of TNFα positive CD8^+^ T cells were detected after stimulation with YopE peptide in both MLN and spleen samples (Fig. 5B left panel). Higher percentages of IFNγ positive CD8^+^ T cells were also observed in spleen samples after treatment with the YopE peptide (Fig. 5C left panel). To better compare the percentage of cytokine-producing cells following peptide stimulation, we normalized the value by taking into consideration the background as measured in the corresponding NS sample and the percentage of tetramer-positive cells. Therefore, the normalized value of 1 indicated that the percentage of cytokine positive cells and the percentage of tetramer positive cells are the same; while higher values suggested an increased sensitivity of the intracellular cytokine staining over tetramers in detecting responding CD8^+^ T cells (Fig. 5BC, right panels). The difference between the OVA- and the YopE-stimulated groups was not significant (Fig. 5BC, right panels). The result indicates that in response to peptide stimulation, the number of CD8^+^ T cells producing either IFNγ or TNFα is proportional to the number of corresponding tetramer positive CD8^+^ T cells present in the sample.

**Fig. 4.**
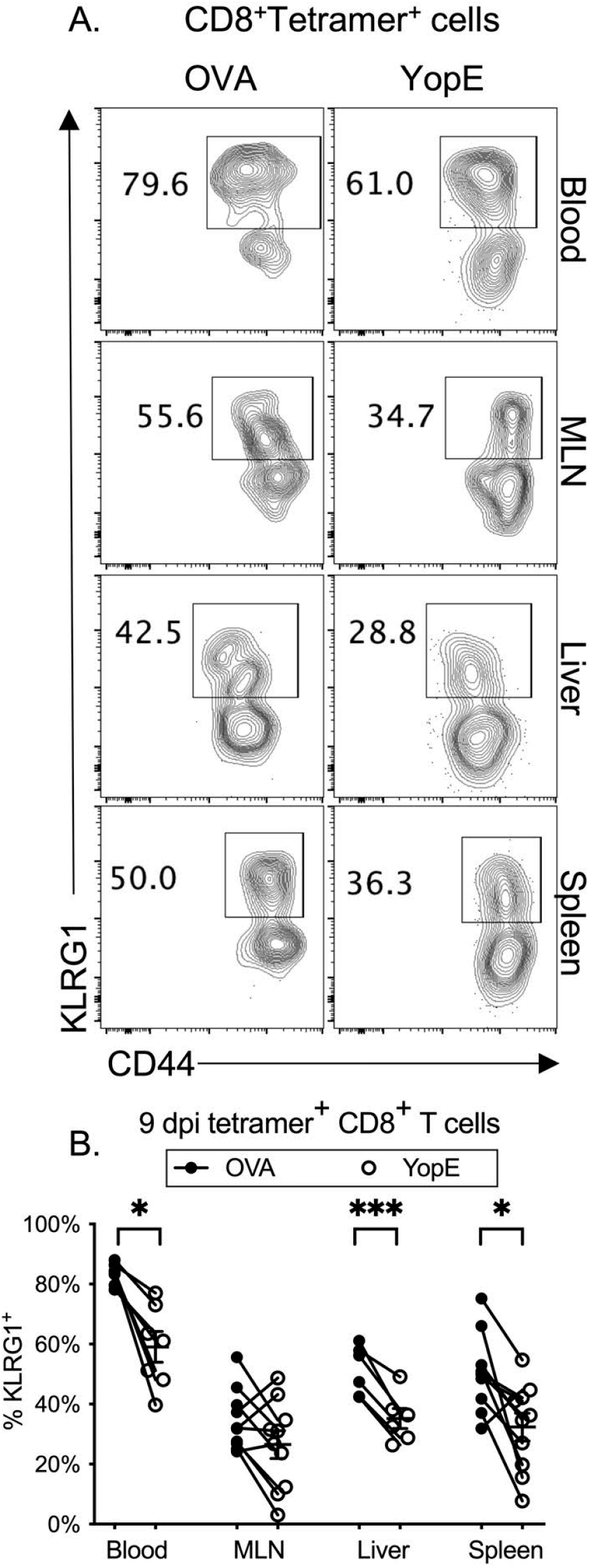
Percentages of OVA-specific or YopE-specific CD8^+^ T cells that are positive for KLRG1 are different at 9 dpi following 32777-OVA infection. C57BL/6 mice were orally infected as in Fig. 1. Percentages of tetramer positive CD8^+^ T cells from the indicated tissues at 9 dpi that are positive for terminal differentiation marker KLRG1 were determined by flow cytometry analysis. A., representative contour plots of CD8^+^tetramer^+^ cells from indicated tissues were further analyzed for CD44 and KLRG1 signal. B., The percentage of KLRG1^high^ cells among OVA- (filled circle) and YopE-specific cells (open circle) plotted according to the tissue sources. Each point represents the value obtained from one mouse and values from the same mouse are connected. The results shown are pooled from 2-4 independent experiments. For blood, *n* equals 7; MLNs, 10; Livers, 6; and spleens, 10. P values were determined with ratio paired t test and values smaller than 0.05 are indicated.

**Fig. 5.**
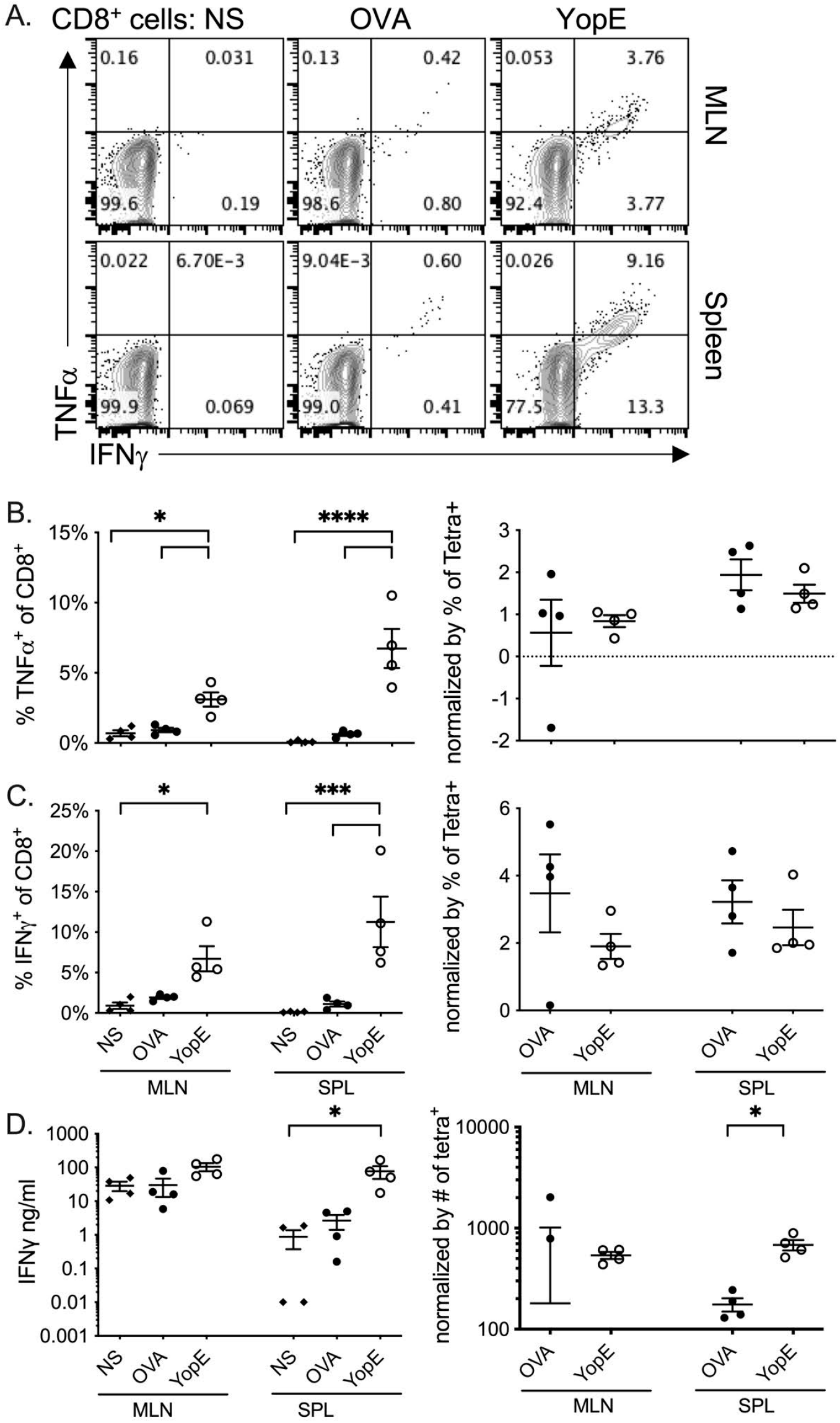
*Ex vivo* production of IFNγ and TNFα after stimulation with OVA_257-264_ and YopE_69-77_ peptide. Mice were orally infected as in Fig. 1. At 9 dpi, total MLN and spleen cells were incubated with peptides YopE_69-77_, OVA_257-264_ or PBS (NS, non-stimulated). (A-C), Intracellular cytokine production from independent wells of cells. Aliquots of cells were incubated as indicated together with BFA for 4.5 h, then stained for indicated intracellular cytokines and surface markers before Flow cytometry. A, representative histographies of CD8^+^ T cells. B and C, percentage of CD8^+^ T cells positive for TNFα (B) or IFNγ (C) were plotted on the left and the values after normalization using the formula (%Cytokine_antigen_-%Cytokine_NS_)/%CD8_antigen_ on the right. In the formula, %Cytokine_antigen or NS_ refers to percent of either TNFα- or IFNγ-positive cells in CD8^+^ T cells following the treatment with either OVA, YopE peptide or PBS, %CD8_antigen_ were percent of OVA or YopE tetramer positive cells in CD8^+^ T cells in the sample. D., Concentrations of secreted IFNγ in the medium of cells incubated with peptide or PBS alone for 48 h without BFA were determined with ELISA and reported as absolute values in ng/ml (left) or after normalization using the formula (C_antigen_ - C_NS_)/Number_antigen_ (right). In the formula, C_antigen or NS_ refers to the concentration of IFNγ in fg/ml following OVA-, YopE-peptide stimulation or NS; and Number_antigen_ refers to the number of antigen specific CD8^+^ T cells in the particular sample (right). Means and standard deviations are indicated. Data shown are combined from two independent experiments and each point represented the value obtained from one mouse <u>and *n* equals 4</u>. P values were determined by Repeated Measure ANOVA, followed by Sidak’s multiple comparisons test. Significant differences were indicated, *, P<0.05, ***, P<0.001, ****, P<0.0001.

To evaluate the abilities of the two types of antigen specific CD8^+^ T cells to secrete IFNγ, aliquots of cells from MLN or spleen of mice infected with 32777-OVA 9 days previously were stimulated *ex vivo* with either YopE or OVA peptides for 48 h or left unstimulated. The concentrations of IFNγ in the medium were then determined. Higher concentrations of IFNγ were observed following stimulation with YopE peptide as compared to the OVA peptide although the differences were not significant. The difference between YopE-stimulated and NS control splenocytes was statistically significant (Fig. 5D, left panel). To compare the cytokine secretion efficiency of an average antigen specific cell, we normalized the concentrations of IFNγ by the number of tetramer positive cells in the sample, which was calculated using results of flow cytometry carried out with an additional aliquot of cells. When background IFNγ concentrations were subtracted, however, two negative values resulted with MLN samples since the IFNγ concentrations were higher in NS than in OVA-stimulated. These two values were not displayed in the log scale in the figure (Fig. 5D, right panel, first column). After normalization, the values obtained from YopE peptide stimulation as compared to OVA peptide were significantly higher in splenocytes but not MLN cells (Fig. 5D, right panel, compare the last two columns). Overall, these results indicate that at 9 dpi YopE-specific and OVA-specific CD8^+^ T cells in spleen and MLN have similar capacities to produce IFNγ or TNFα in response to peptide stimulation *ex vivo*, although YopE-specific CD8^+^ T cells in spleen may have a subtle advantage in secretion of IFNγ under these conditions.

These results prompted us to further evaluate a potential difference in the effector differentiation of these two populations of antigen-specific cells over a longer time course. Following Lm infection, through differential expression of CD127 and KLRG1, antigen specific CD8^+^ T cells can be divided into short-lived effector cells (SLECs, CD127^neg^KLRG1^+^), double positive effector cells (DPECs, CD127^+^KLRG1^+^), memory precursor effector cells (MPECs, CD127^+^KLRG1^neg^) and early effector cells (EECs, CD127^neg^KLRG1^neg^ (41). SLECs undergo apoptosis during T cell contraction, while MPECs form long-lived memory cells. A similar process of memory formation has yet to be established following *Yersinia* infection, but a distinction in SLEC, DPEC or MPEC populations among OVA- and YopE-specific cells would be consistent with a difference between these two populations of antigen-specific CD8^+^ T cells in differentiation. In either livers or spleens, sufficient numbers of OVA-specific cells were identified that warranted such analysis. For both OVA- and YopE-specific CD8^+^ T cells, at 9 dpi, the percentages of all four effector populations displayed a wide range (Fig. 6). In average, EECs were the largest population at this time point for both tissues (Fig. 6BC). In spleens, consistent with a later peak time (Fig. 2E, bottom panel), the YopE-specific CD8^+^ T cells contained higher percentages of EECs than those OVA-specific cells (Fig. 6C). With progress of infection, the ranges in the percentages of these effector populations decreased (Fig. 6BC). Higher levels of SLECs and lower levels of MPECs were identified from YopE-specific as compared to OVA-specific CD8^+^ T cells and the differences were significant in both livers and spleens at 14 and 30 dpi (Fig. 6B and C). These observations are consistent with a potential difference in memory formation of YopE- and OVA-specific CD8^+^ T cells.

**Fig. 6.**
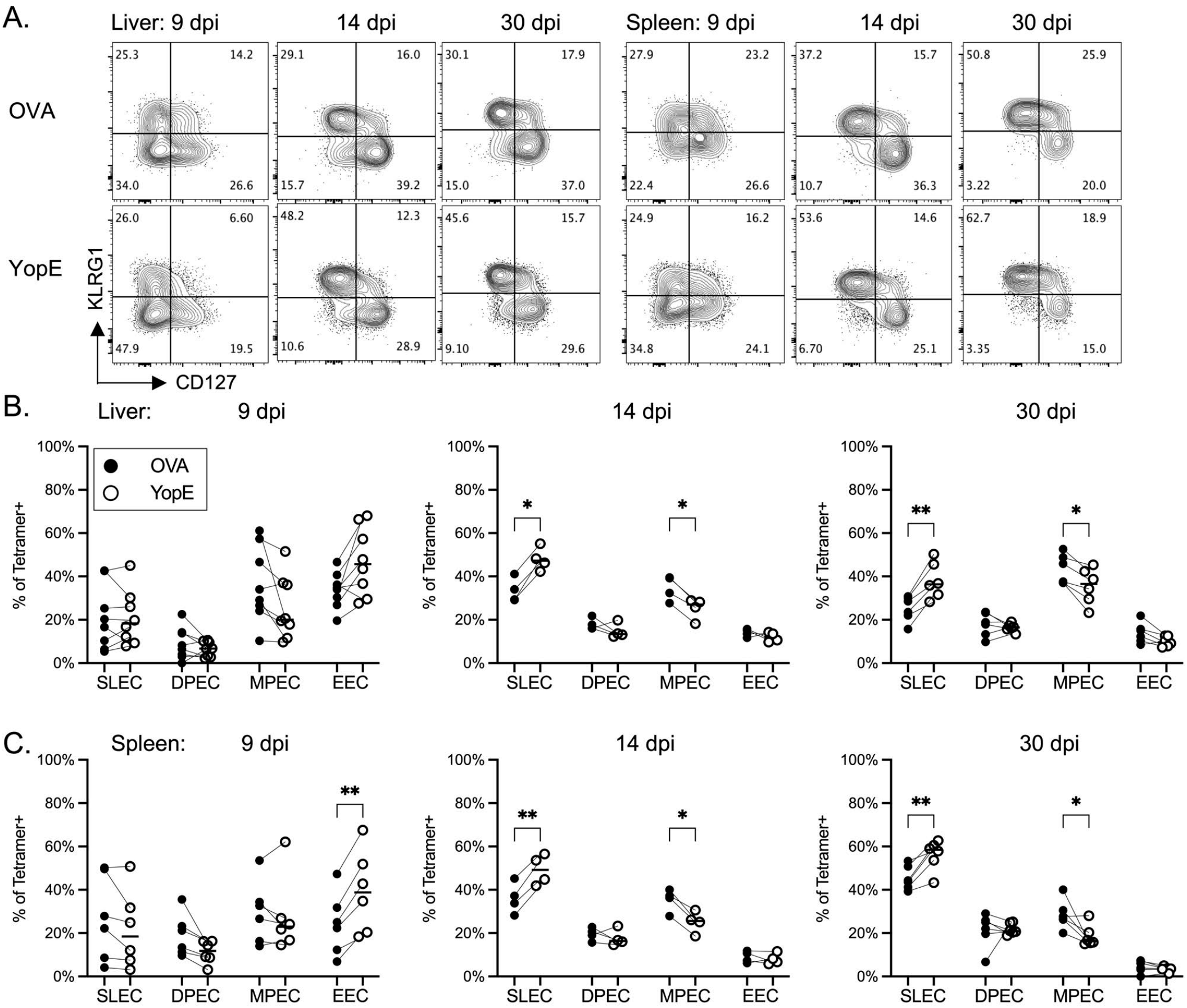
The OVA and YopE-specific CD8^+^ T cells differ in the composition of SLEC and MPEC populations defined by CD127 and KLRG1 expression following infection with 32777-OVA. Expression of CD127 and KLRG1 in alive tetramer positive CD45^+^TCRβ^+^CD8α^+^CD44^hi^ T cells was determined at 9, 14 and 30 dpi. A., representative contour plots of OVA-specific (top) and YopE-specific (bottom) CD8^+^ T cells from liver (left) and spleen (right) at indicated days post infection. The percentages of SLEC (CD127^neg^KLRG1^+^), DPEC (CD127^+^KLRG1^+^) ^+^), MPEC (CD127^+^KLRG1^neg^) and EEC (CD127^neg^KLRG1^neg^) in OVA-specific (filled circle) and YopE-specific (open circle) cells in livers (B) and spleens (C) plotted according to the dpi indicated. Values obtained from the same animal were connected. Data shown are summary from two experiments for 9 and 30 dpi and one for 14 dpi. With livers, *n* equals 8, 4, and 6 for 9, 14 and 30 dpi, respectively; spleens, 6, 4 and 6 for 9, 14 and 30 dpi, respectively. P values were determined with Repeated measures ANOVA followed with Sidak’s multiple comparisons, and those smaller than 0.05 were indicated.

### Higher levels of intestinal YopE-specific compared to OVA-specific CD8^+^ T cells following 32777-OVA infection

To evaluate the antigen specific CD8^+^ T cell response in intestinal tissue, lymphocytes were isolated and analyzed with flow cytometry after staining. Similar to MLNs, livers and spleens, the percentages of a wide range of distribution were observed in the levels of YopE-specific CD8^+^ T cells among intestine intraepithelial lymphocytes (IEL) or the lymphocytes from lamina propria (LP) at 9 dpi (Fig. 7ABC). From 7 dpi to 30 dpi, the mean percentage of CD8^+^ T cells positive for YopE-tetramer in IEL remained relatively stable, while the values for OVA-specific cells decreased over this time period (Fig. 7B). Similar results were observed in the LP, although the percentages of YopE-specific CD8^+^ T cells peaked 14 dpi (Fig. 7C). Additionally, the ratio of YopE-specific cells over OVA-specific CD8^+^ T cells also increased from 9 dpi to 30 dpi, a difference that was significant in the LP (Fig. 7DE).

**Fig. 7.**
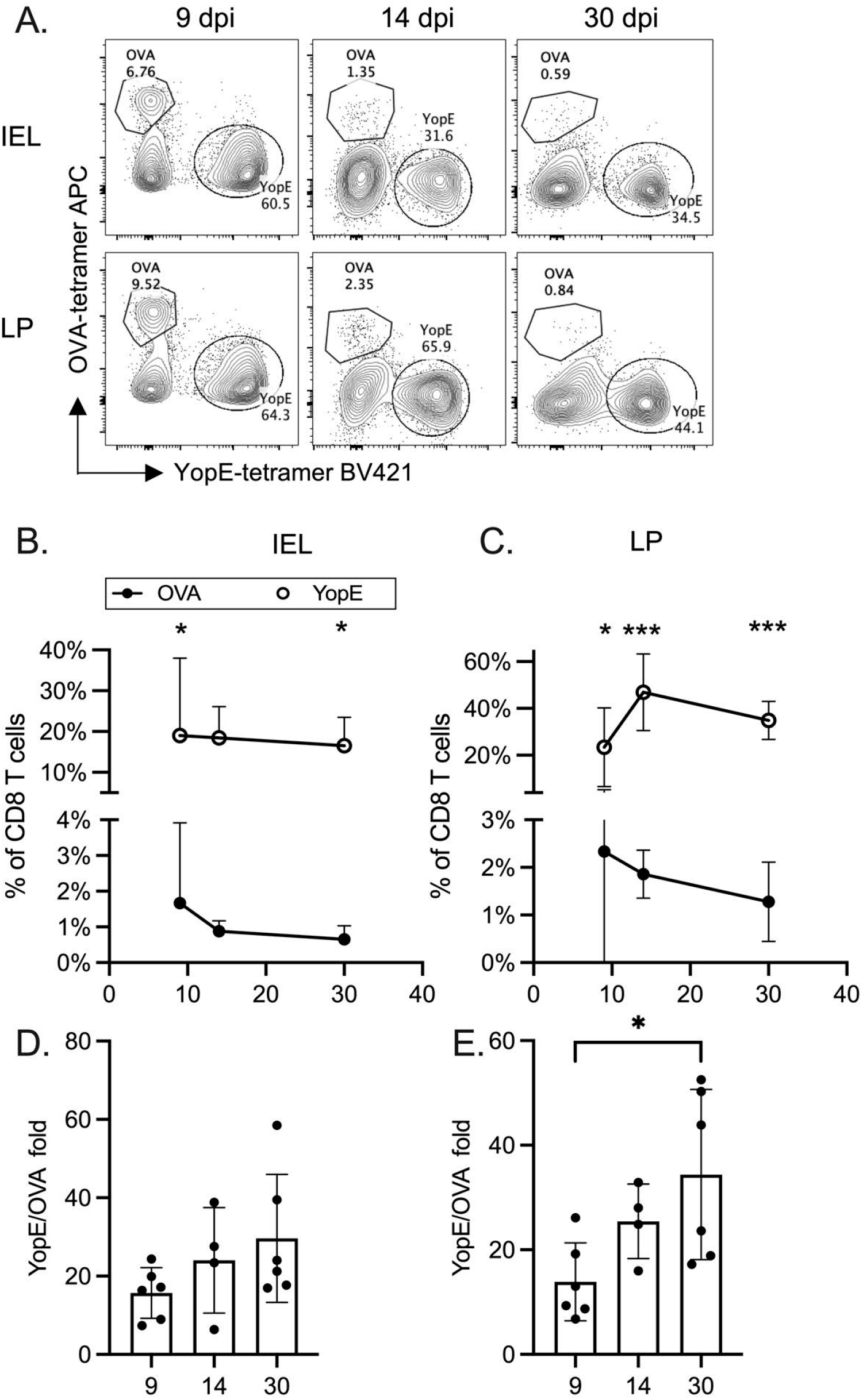
Different levels of OVA-specific and YopE-specific CD8^+^ T cells in IEL and LP. Tetramer positive cells were identified from IEL and LP using flow cytometry analysis following infection. A., representative contour plots of alive CD45^+^TCRβ^+^CD8α^+^CD44^hi^ cells from IEL (top) and LP (bottom) for tetramer signals at indicated dpi. B-C. The percentages of OVA-tetramer^+^ (filled circle) and YopE-tetramer^+^ (open circle) in CD8^+^ cells (CD45^+^TCRβ^+^CD8α^+^) from either IEL (B) or LP (C) were plotted according to the dpi indicated. P values were determined with Two-way ANOVA (Mixed-effects model) followed by Sidak’s multiple comparisons test and values smaller than 0.05 are indicated. D-E., the ratio of YopE-specific CD8^+^ T cells over OVA-specific ones from either IEL (D) or LP (E) according to the dpi indicated. P values were determined with Brown-Forsythe ANOVA test followed with Dunnett’s T3 multiple comparisons test and values smaller than 0.05 are indicated. Data shown are summary from one (14 dpi) or two experiments (rest) and *n* equals 6, 4 and 6 for 9, 14 and 30 dpi, respectively.

To evaluate memory formation, the antigen-specific CD8^+^ T cells were quantified for CD103 and CD69 expression. Bacterial or viral infection results in CD8^+^ T cell effector activation and trafficking into peripheral tissues such as skin, lung or intestine to differentiate into T_RM_ cells to provide accelerated pathogen clearance upon recounter (42). Following oral *Y. pseudotuberculosis* infection, two populations of T_RM_ cells have been identified—the classical CD69^+^CD103^+^ T_RM_ cells and the CD69^+^CD103^neg^ cells found in LP (37, 43). Among IELs, the classical CD69^+^CD103^+^ T_RM_ cells were identified from both OVA-specific and YopE-specific CD8^+^ T cells (Fig. 8AB). As reported before (37), the percentages of CD69^+^CD103^+^ T_RM_ cells increased from 9 to 30 dpi among both OVA-specific and YopE-specific CD8^+^ T cells (Fig. 8B, left panel). Lower percentages of YopE-specific as compared to OVA-specific CD8^+^ T cells were observed in the these CD69^+^CD103^+^ T_RM_ cell populations throughout the time course analyzed, and this difference was significant at 30dpi (Fig. 8B, left panel). In LP, although both CD69^+^CD103^+^ and CD69^+^CD103^neg^ T_RM_ cells were identified, their percentages were variable among the two populations of antigen-specific CD8^+^ T cells over time (Fig. 8AB). In the LP there was a significantly lower percentage of YopE-specific as compared to OVA-specific CD8^+^ T cells among the CD69^+^CD103^+^ T_RM_ cell population at 9dpi and the CD69^+^CD103^neg^ T_RM_ cell population at 30 dpi (Fig. 8B, middle and right panels, respectively). Therefore, overall, these results show that there are higher numbers of intestinal YopE-specific compared to OVA-specific CD8^+^ T cells at 9dpi following infection with 32777-OVA, and the former cell type dominance increases between 9 and 30dpi. In addition, both types of antigen-specific cells can become T_RMs_, although YopE-specific CD8^+^ T cells may acquire these phenotypes less efficiently during the time analyzed.

**Fig. 8.**
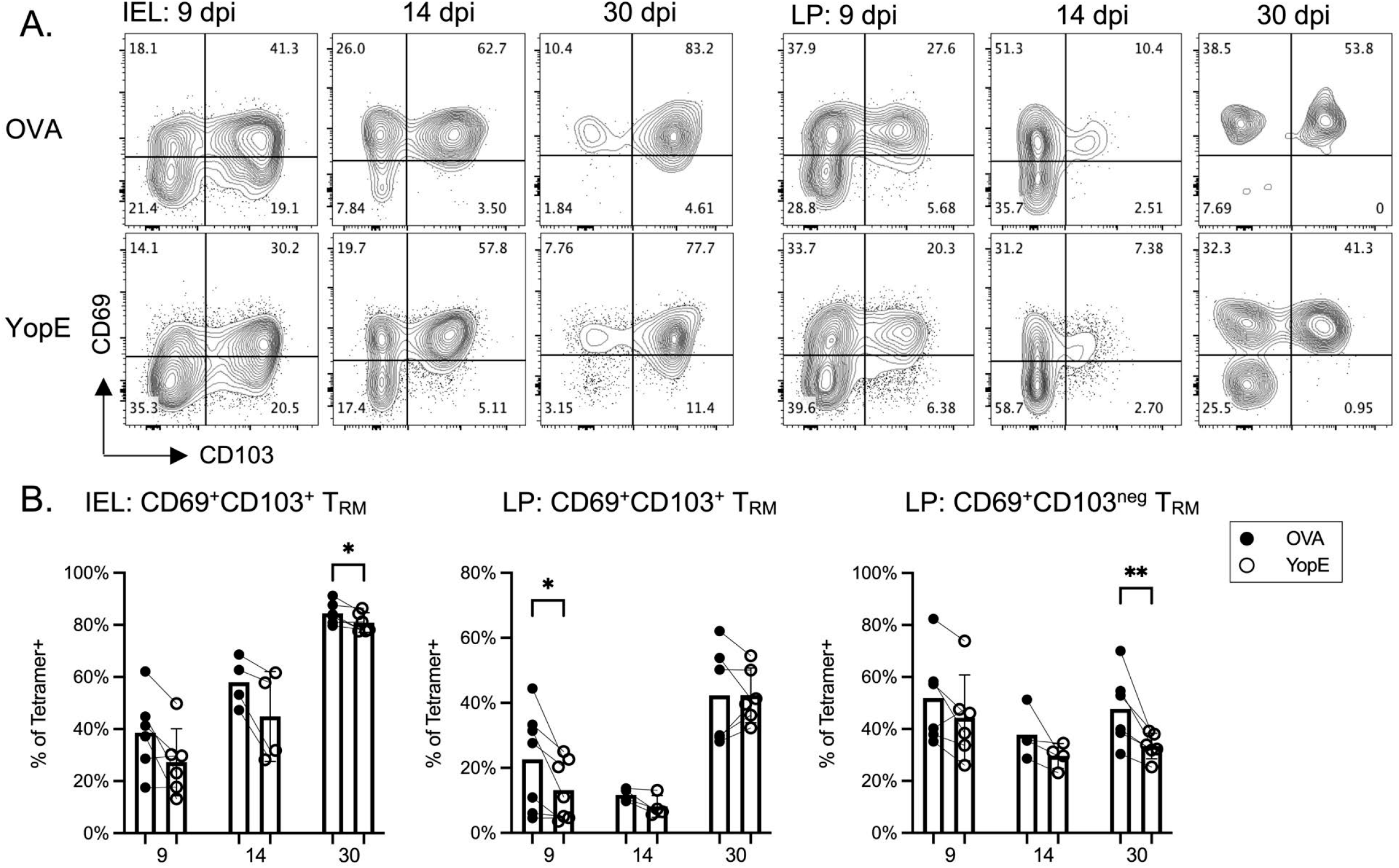
T_RM_ formation among OVA-specific and YopE-specific CD8^+^ T cells identified from IEL and LP following 32777-OVA infection. Tetramer positive cells identified from IEL and LP were analyzed for expression of CD103 and CD69 using flow cytometry analysis. A., representative contour plots of either OVA-tetramer^+^ (top) or YopE-tetramer^+^ (bottom) cells isolated from IEL (left) or LP (right) for CD103 and CD69 signals at indicated dpi. B. The percentages of indicated types of T_RM_ cells among OVA-tetramer^+^ (filled circle) and YopE-tetramer^+^ (open circle) cells were plotted according to the dpi indicated. Data shown are summary from one (14 dpi) or two experiments (rest) and *n* equals 6, 4 and 6 for 9, 14 and 30 dpi, respectively. P values were calculated with ratio paired t test and values smaller than 0.05 are indicated.

## Discussion

We set out to characterize the antigen specific CD8^+^ T cell response in foodborne *Y. pseudotuberculosis* infection and expressed the model CD8^+^ T cell epitope OVA_257-264_ in the same protein as the native *Yersinia* epitope YopE_69-77_. It was initially a surprise that the YopE-specific CD8^+^ T cell response was several times higher than the OVA-specific response. Here we showed that the YopE-specific CD8^+^ T cells have a high precursor frequency in naïve C57BL/6 mice and that this high precursor frequency is associated with the large and relatively more sustained YopE-specific CD8^+^ T cell response than that of OVA-specific cells following *Y. pseudotuberculosis* infection. In addition, we provide evidence suggesting that these two types of antigen specific CD8^+^ T cells may be different in additional aspects including effector differentiation, the ability to secrete cytokines <u>and memory formation</u>.

The number of T cell precursors specific for different antigens in an individual can vary greatly (reviewed in (40)). Using tetramer based cell enrichment, it was determined that the number of naïve antigen specific CD8^+^ T cells ranged from ~15-1500 cells per mouse for the 25 different viral antigenic epitopes studied and the model antigen OVA_257-264_ (40). Two independent studies identified that the number of CD8^+^ precursors specific for murine cytomegalovirus M45:D^b^ is the highest, either as 1500 cells/mouse or at 603 cells/mouse but 4.6 times as abundant as that of OVA-specific precursors (39, 44). Our results here indicated that the average number of YopE-specific CD8^+^ precursor is 1387/mouse or 4.6 times as abundant as that of OVA-specific ones (Fig. 3C). Even considering the differences in measurement conducted at different laboratories, this result indicates that the YopE-specific CD8^+^ T cell precursors are among the most abundant precursors known.

The magnitude of a primary T cell response is influenced by the amount and duration of the antigenic peptide presented on MHC molecules of APCs in secondary lymphoid organs. Recently, with the quantification of T precursors, the frequency of the naïve precursors was also recognized as an important factor deciding the size of a primary response (40). Here, when the two epitopes of YopE_69-77_ and OVA_257-264_ were co-expressed in the same protein, theoretically they would be processed and presented equally. In addition, both types of antigen specific cells are exposed to the same cytokine environment, which impact T cell activation (4). Therefore, it was initially surprising when we first observed that the OVA-specific CD8^+^ T cell response was much smaller than that of the YopE-specific response (Fig. 2), especially since the OVA epitope has a slightly stronger calculated affinity for MHC I than the YopE epitope, IC50 of 17 nM and 20 nM respectively (24). In this regard, the large number of YopE-specific CD8^+^ T cell precursors in naive C57BL/6 mice correlates well with our observation. Previously, Bergsbaken et al. had adoptively transferred 10^4^ OT-I T cells to mice before infection with *Y. pseudotuberculosis* expressing a YopE-ovalbumin fusion protein (37). This essentially boosted the OVA-specific CD8^+^ T cell “precursor” levels to more than that of native YopE-specific cells. However, the OT-I T cell response they observed was still several times smaller than the YopE-specific response following infection with the OVA-expressing *Y. pseudotuberculosis* strain. They reasoned that their YopE-ovalbumin protein was expressed from a weaker promoter, while the endogenous YopE was expressed at much higher levels. As a result, a smaller number of OVA peptides may have been presented on MHC I molecules. Consequently, the smaller OVA-response was due to the smaller antigen load of OVA as compared to YopE (37). Together, their study and ours presented here indicate that different factors, either the abundance of the antigenic peptides presented, or the abundance of the precursors, could differentially impact the magnitude of the primary CD8^+^ T response to even the same kind of pathogen.

Besides the abundance of precursors, the affinity of TCR to pMHC complexes has also been implicated to influence the primary virus-specific cytotoxic T cell response (reviewed in (45)). The affinity of TCR to pMHC may dictate the expansion of individual CD8^+^ T cell clones to result in the number of progenies varying over three orders of magnitude as observed in individual T cell transfer experiments (14). Alternatively, the local environment impacts the differential expansion of T cells. Our results of increased ratio between YopE/OVA-specific CD8^+^ T cells with the progress of infection from 7 dpi to 30 dpi in multiple tissues including intestines are consistent with differential expansion of T cell clones (Fig. 2 F-H and Fig. 7DE)—shorter division time of the YopE-specific CD8^+^ T cells resulting in faster expansion. In MLN, livers and spleens, the YopE/OVA T cell ratios at 7 dpi are close to the ratio of YopE/OVA precursor measured in naïve mice. Then through expansion of random combinations of both YopE and OVA-specific T cell clones, the values of this ratio became variable, but in average, the ratio increased overtime. Division speed of antigen-specific CD8^+^ T cells has been shown to impact central memory formation in that central memory precursors divide slower (16). This is consistent with our observation that higher percentage of OVA-specific cells are MPECs at 14 and 30 dpi in both livers and spleens (Fig. 6). Furthermore, when T_RM_ cells were quantified from IELs or the lymphocytes from LP, higher percentages of OVA-specific cells were found to bear T_RM_ markers at 30dpi (Fig. 8B). In addition, our results suggested that an average YopE-specific CD8^+^ T cell effector may be more efficient in secretion of cytokines such as IFNγ than an OVA-specific cell. Following *ex vivo* stimulation with either OVA or YopE antigenic peptide, more IFNγ was secreted per responding cell from the spleen but not MLN (Fig. 5D).

How could the lower frequency of the OVA-specific precursors associate with general lower TCR to pMHC avidity, and therefore slower division speed during activation? It was shown recently that CD4^+^ T cells cross reacting with self-peptides may be deleted during development and small naïve T cell populations experience extensive clonal deletion (46). Furthermore, the T cells that survived clonal deletion express TCRs with lower affinity (46). A similar process has yet to be demonstrated for CD8^+^ T cells, although it is plausible that the cells from a smaller precursor population also express TCRs with lower affinities than those from a larger precursor population. Although speculative, the possibility that antigen specific CD8^+^ T cells from a rare precursor population express TCR with lower affinity merits future investigation.

In summary, our finding that there is a high precursor frequency of YopE-specific CD8^+^ T cells partially explains the longstanding mystery behind the exceptional dominance of the protective YopE_69-77_ epitope in C57BL/6 mice (21–26). The high precursor frequency of YopE-specific CD8^+^ T cells may contribute to the increased dominance of these cells over OVA-specific cells during the course of 32777-OVA infection, as wells as differences in effector and memory populations of these two cell types. These results have important implications for studying mechanisms of protection afforded by YopE-specific CD8^+^ T cells in *Yersinia* infection models with C57BL/6 mice. In addition, our findings suggest that 32777-OVA infection of C57BL/6 mice will provide a useful system for better understanding how precursor frequency impacts divergent CD8^+^ T cells responses in general.

## Materials and Methods

### Bacterial Strains

The *Y. pseudotuberculosis* strains used in this study are serogroup O:1 strain 32777 and its derivative 32777-OVA. To generate strain 32777-OVA, oligos 5’-ccggctAGcATAATCAACTTTGAAAAACTG-3’ and 5’-CCGGCAGTTTTTCAAAGTTGATTATGCTag-3’ were synthesized and annealed at room temperature, then inserted into the AgeI site of plasmid pSB890-YopEplus (25). The resulting plasmid pSB890-YopEOVA was used in standard allelic exchange procedures to place the SIINFEKL epitope in the linker region of YopE coding sequence after residue 85 of YopE (Fig. 1A) in the virulence plasmid together with an NheI site to facilitate identification.

### Infection of mice

C57BL/6 mice were purchased from Jackson Laboratory and were used within 8 to 16 weeks of age. For foodborne infection, overnight *Y. pseudotuberculosis* culture grown in Luria-Bertani (LB) medium at 28°C was washed once and resuspended in phosphate-buffered saline (PBS) to achieve the desired CFU/ml. For each mouse to be infected, a 50 μL volume of the suspension was applied to single ~0.5 cm^3^ piece of white bread placed on a thin layer of bedding in the bottom of a fresh cage. One mouse was introduced into the cage. The entire cage was kept in the dark and checked periodically until the bread was consumed, usually within 1 to 2 hours. Then the mouse was returned to its original cage.

### Processing of tissues

At indicated days post infection or when death was imminent, mice were euthanized by CO_2_ asphyxiation. Blood was collected through heart punctual, and red blood cells were lysed. Mouse MLNs, spleens and livers were dissected and cut into two pieces aseptically and weighed. Half of spleens and MLNs were homogenized with a 3 ml syringe plunger in 5 ml of collection medium (RPMI-1640 containing 5% Bovine serum and 20 U/ml Penicillin, 20 μg/ml Streptomycin and 5 ng/ml Gentamicin). Half of the livers were processed to enrich lymphocytes according to manufacturer’s instructions with a gentleMACS Dissociator in C tubes (Miltenyi Biotec). Alternatively, liver pieces were treated with collagenase at 100 units/ml in collagenase buffer (RPMI-1640 medium containing 10% Fetal Bovine Serum, 1 mM CaCl_2_, 1 mM MgCl_2_, 20 U/ml Penicillin, 20 μg/ml Streptomycin and 5 ng/ml Gentamicin) at 37°C for 40 minutes. The homogenates were then meshed through 70 μm conical filter cap to obtain single cell suspension, and lymphocytes were enriched with a Percoll gradient. Small intestine intraepithelial lymphocytes (IELs) and lamina propria (LP) lymphocytes were collected as previously described (34, 35). Briefly, small intestine was cut open and sliced into short pieces after removing attached fat, mesentery, Peyer’s patches and mucous. IELs were collected following two treatments of DTE buffer (1X HBSS containing 1 mM HEPES, 2.5 mM NaHCO_3_, and 1 mM dithioerythritol) at 37°C for 20 minutes each. Then after removal of epithelial layers with EDTA buffer, LP lymphocytes were released after treatment with collagenase at 100 units/ml for 40 minutes at 37°C. IELs and LP lymphocytes were further enriched with a Percoll gradient.

The MLNs, spleens or the other half of livers were homogenized in FACS buffer (PBS containing 0.2% Bovine Serum Albumin and 2 mM EDTA). Aliquots from the homogenized tissues were serially diluted in LB and plated (100 μl) on LB agar to determine bacterial colonization by CFU assay, and the limit of detection was 100 CFU or a log10 CFU value of 2. All animal procedures were approved by the Stony Brook University Institutional Animal Care and Use Committee.

### Flow Cytometry

Viable cells from the single cell suspension or enriched lymphocytes were counted using trypan blue exclusion with Vi CELL XR cell Viability Analyzer (Beckman Coulter). Initially, suspended cells (1X10^6^) were blocked using anti-mouse CD16/CD32 (FcgIII/II receptor) clone 2.4G2 (BD) and labeled with allophycocyanin (APC)-conjugated MHC class I tetramer K^b^YopE_69-77_, which was provided by the NIH Tetramer Core Facility (Emory University, Atlanta, GA), or OVA_257-264_, at room temperature for 1 h and fluorophore-conjugated antibodies on ice for 20 minutes. The antibodies used were AlexaFluor488 or PE anti-mouse CD8α (53-6.7, BD), and PE/Cy7 anti-mouse CD3e (clone 145-2C11, BD), Brilliant Violet 421™ or PE anti-CD44, Brilliant Violet 421™ or PerCP/Cy5.5 anti-mouse CD4 (RM4-4), PE anti-mouse/human KLRG1 (2F1/KLRG1), V500 anti-mouse CD45.2 (104, BD). CD8^+^ T cells were gated as CD3^+^CD8^+^ events, or alive CD45.2^+^CD3^+^CD8^+^ population whenever possible. Labeled cells were analyzed using a Cytek DXP 8 color upgrade. To analyze memory formation, up to 5X10^6^ cells were mixed with anti-mouse CD16/CD32 and stained with Brilliant Violet 510™ anti-CD45, PerCP/Cy5.5 anti-TCRβ, Brilliant Violet 786™ anti-CD8α, APC-eF780 anti-CD44, APC-conjugated OVA-tetramer, Brilliant Violet 421™-conjugated YopE-tetramer, PE anti-CD127, FITC anti-KLRG1, PE/Dazzle 594 anti-CD103 and PE/Cy7 anti-CD69. Labeled cells were analyzed using a FACSDiva. Antibodies were from BioLegend unless indicated otherwise. Data were analyzed with FlowJo software (Tree Star).

### Secretion of IFNγ and intracellular cytokine staining following *ex vivo* stimulation

To stain for intracellular cytokine, an aliquot of suspended splenocytes or cells from MLN (1X10^6^ cells) were incubated in 200 μL of complete T cell medium (Dulbecco modified Eagle medium supplemented with 10% heat-inactivated fetal bovine serum, 12.5 mM HEPES, 2 mM L-glutamine, 1 mM sodium pyruvate, 1 mM penicillin-streptomycin and 55 μM β-mercaptoethanol) containing 10 nM either OVA_257-264_ (H_2_N-SIINFEKL-OH) or YopE_69-77_ peptide and brefeldin A (BFA, Sigma, 5 μg/ml) for 4.5 h at 37°C. Then the cells were washed and stained first for CD8α, CD4, CD3e and CD45.2 as described above, then the cells were fixed and permeabilized with BD Cytofix/Cytoperm kit according to manufacturer’s instructions. Finally, the cells were stained with PE anti-mouse IFNγ (clone XMG1.2) and PerCP/Cy5.5 anti-mouse TNFα (clone MP6-XT22, both from BioLegend). Isotype-matched antibodies were used to control for nonspecific binding. To determine the secretion of IFNγ, cells were incubated for 48 h without BFA. Then the concentrations of IFNγ in the supernatant were determined by enzyme-linked immunosorbent assay (ELISA) following the manufacturer’s instructions (R&D systems, Inc.).

### Precursor determination

Naïve antigen-specific CD8^+^ T cells were determined using the MHCII Tetramer Pulldown Protocol (36) with modifications. In brief, the spleen, inguinal, axillary, brachial, submandibular, cervical, para-aortic and mesenteric lymph nodes were dissected, then single cell suspensions in sorter buffer (PBS with 2% fetal bovine serum) were filtered through a 70 μm nylon Cell Strainer and pooled, spun down and re-suspended in sorter buffer containing 2% mouse serum and 8 μg Fc blocking antibody 2.4G2 in a final volume about twice that of the pellet. Then antigen specific cells were enriched after sequential incubation with either OVA- or YopE-specific tetramers conjugated with APC then Miltenyi anti-APC microbeads, followed by standard LS column separation. Eluted cells were further incubated with OVA-specific PE-tetramer or YopE-specific BV421-tetramer, then CD8α-PerCP/Cy5, CD90.2-PE/Cy7 together with FITC-conjugated dump antibodies (B220, Gr1, CD11c, F4/80 and CD4). Dead cells were disregarded based on staining with Alexa Fluor 700 Carboxylic acid, succinimidyl Ester. AccuCheck counting beads from Invitrogen were included in the samples of stained cells, and the total number of cells in each sample was determined with the formula: Cell count/bead count x bead stock concentration x bead volume/cell volume x total sample volume.

### Statistical Analysis

Statistical analyses were performed in Prism (GraphPad Software). Tests used were listed in each figure. The p values smaller than 0.05 were considered significant and were indicated as follows: *, p<0.05; **, p<0.01; ***, p<0.001; and ****, p<0.0001. Flow data were exported from FlowJo to Microsoft Excel for initial calculation.

## Acknowledgements

The authors thank the NIH Tetramer Core Facility for providing tetramer reagents, and Josh Obar, Adrianus W. M. van der Velden and Camille Khairallah for valuable comments. This research was supported by an Institutional Research and Academic Career Development Award (K12-GM-102778 to ZQ) and a National Institute of Allergy and Infectious Diseases of the National Institutes of Health grant (R01AI099222 to JBB).

